# Broad applicability of a streamlined Ethyl Cinnamate-based clearing procedure

**DOI:** 10.1101/346247

**Authors:** Wouter Masselink, Daniel Reumann, Prayag Murawala, Pawel Pasierbek, Yuka Taniguchi, Jürgen A. Knoblich, Elly M. Tanaka

## Abstract

Turbidity and opaqueness are inherent properties of tissues which limit the capacity to acquire microscopic images through large tissues. Creating a uniform refractive index, known as tissue clearing, overcomes most of these issues. These methods have enabled researchers to image large and complex 3D structures with unprecedented depth and resolution. However, tissue clearing has been adopted to a limited extent due to a combination of cost, time, complexity of existing methods and potential negative impact on fluorescence signal. Here we describe 2Eci (2^nd^ generation Ethyl cinnamate based clearing method) which can be used to clear a wide range of tissues, including cerebral organoids, *Drosophila melanogaster,* zebrafish, axolotl, and *Xenopus laevis* in as little as 1-5 days while preserving a broad range of fluorescence proteins including GFP, mCherry, Brainbow, and alexa-fluorophores. Ethyl cinnamate is non-toxic and can easily be used in multi-user microscope facilities. This method will open up clearing to a much broader group of researchers, due to its broad applicability, ease of use, and non-toxic nature of Ethyl cinnamate.

**Summary statement:** The non-toxic, broadly applicable, and simplified protocol of 2Eci tissue clearing makes it possible for non-specialist labs to use clearing approaches on conventional inverted microscopes.

## Introduction

Methods to optically clear tissues using refractive index matching have been transformative for imaging large, 3-dimensional tissues. Such methods have allowed long-distance mapping of axonal projections and reconstruction of entire embryos (Belle *et al*., 2014; Economo *et al*., 2016; Belle *et al.,* 2017). Despite the importance of these methods, the widespread, daily use of clearing agents to quantify cell populations in whole-mount preparations has seen limited use in rapidly evolving fields such as developmental biology, organoid research or regeneration biology due to cumbersome aspects associated with each method. Aqueous-based clearing media such as Clarity and SeeDB, require long incubation times for equilibration and immunostaining typically requiring days to weeks to complete depending on tissue size (see Table 1). This becomes prohibitive for rapidly screening different experimental conditions. Organic-solvent based methods bypass long incubations times due to extraction of lipids and other organic material in the sample, yet are often either toxic, show limited clearing or reduced preservation of fluorescence protein signal (See Table 1). We aimed to overcome these shortcomings to produce a rapid, yet effective and non-toxic clearing protocol that preserves fluorescent protein/antibody signal. Such a method would allow, for example, the use of whole-mount organoid imaging for genetic or chemical screening. The method would also allow for interrogation of fluorescent protein-expressing transgenic animals combined with immunofluorescence to quantify/characterize discrete populations in complex samples such as an adult fly or regenerating axolotl limbs. Here we describe the combination of sample dehydration in 1-propanol_pH9_ followed by refractive index matching with the organic compound ethyl cinnamate (Ethyl 3-phenyl-2-propenoate) as an ideal protocol for rapid, non-toxic sample preparation that preserves protein and labeled-antibody fluorescence which we call 2Eci (2^nd^ generation Ethyl cinnamate based clearing method). We apply the 2Eci method to cerebral organoid characterization, whole-animal and whole-appendage imaging.

**Table 1:** overview of various commonly used clearing methods.

| Method | Aqueous/Organic | Timescale | Toxicity? | Clearing | GFP preservation | reference |
| --- | --- | --- | --- | --- | --- | --- |
| SeeDB | A | 3 days | no | ++ | ++++ | (Ke, Fujimoto and Imai, 2013) |
| Clarity | A | 8 days | no | +++ | ++++ | (Chung *et al.*, 2013) |
| FluoBABB | O | >2 days | yes | ++ | +++ | (Schwarz *et al.*, 2015) |
| Eci/ETOH | O | >5 days | no | +++ | + | (Klingberg *et al.*, 2017) |
| Eci/Prop<sub>pH9</sub> | O | 25 hours | no | +++ | +++ | Current manuscript |

## Results and Discussion

### Establishment of clearing conditions

Our aim was to develop a rapid clearing protocol that preserves GFP fluorescence. We therefore focused on organic-chemical based protocols for their rapidity, and aimed to optimize efficiency of clearing and preservation of fluorescence signal. We assessed clearing efficiency and preservation of GFP fluorescence using cerebral organoids sparsely labeled with a population of CAG:GFP^+^ expressing cells. Human cerebral organoids are a powerful 3D culture system that reconstitutes the early development of discrete brain regions(Lancaster *et al.,* 2013). These organoids provide a reductionist approach to understand aspects of human brain development in-vitro (Bagley *et al.,* 2017). Uncleared cerebral organoids are highly turbid (Fig. 1A). While FluoClearBABB (Fig. 1B) provides moderate improvement in turbidity, ethanol dehydration followed by refractive index matching using ethyl cinnamate as previously described (Klingberg *et al.,* 2017) cleared cerebral organoids (Fig. 1C). However, GFP fluorescence intensity, while still present, was significantly reduced resulting in the loss of ability to detect detailed cellular morphology (Fig. 1E). Based on reports that dehydration using alcohols adjusted to alkaline pH levels can preserve GFP fluorescence (Schwarz *et al.,* 2015) we assessed clearing efficiency and GFP preservation in a series of alcohols adjusted to pH9 (Fig. 1D-H). We found that dehydration using methanol_ph9_ and ethanol_ph9_, reduced fluorescent signal to approximately 1% and 5% of uncleared signal, so that morphological details of GFP^+^ cells could no longer be observed (Fig. 1D-F, I). Cerebral organoids dehydrated with either 4-butanol _ph9_ or 1-propanol _ph9_ displayed a higher intensity of GFP signal at approximately 50% and 75% of uncleared signal (Fig. 1G-H, I). To asses clearing efficiency, we examined imaging depth independent of GFP fluorescence by recording auto-fluorescence levels at 488 nm wavelength. We found that total autofluorescence levels are comparable across dehydrating agents although increased compared to unfixed control samples (Fig. S1A). Methanol _ph9_, ethanol _ph9_, and 1-propanol _ph9_ allow for autofluorescence recordings through the whole organoid (>1400 μm), while 4-butanol _ph9_ mediated clearing yielded only 500 μm penetration into the organoid (Fig. 1J). We conclude that the combination of 1-propanol _ph9_-mediated dehydration followed by Ethyl cinnamate mediated refractive index matching allows for efficient clearing while preserving sufficient levels of GFP and can be completed in as little as 25 hours (Fig. 1K-M). We call this method 2Eci (2^nd^ generation Ethyl cinnamate mediated clearing).

**Figure 1:**
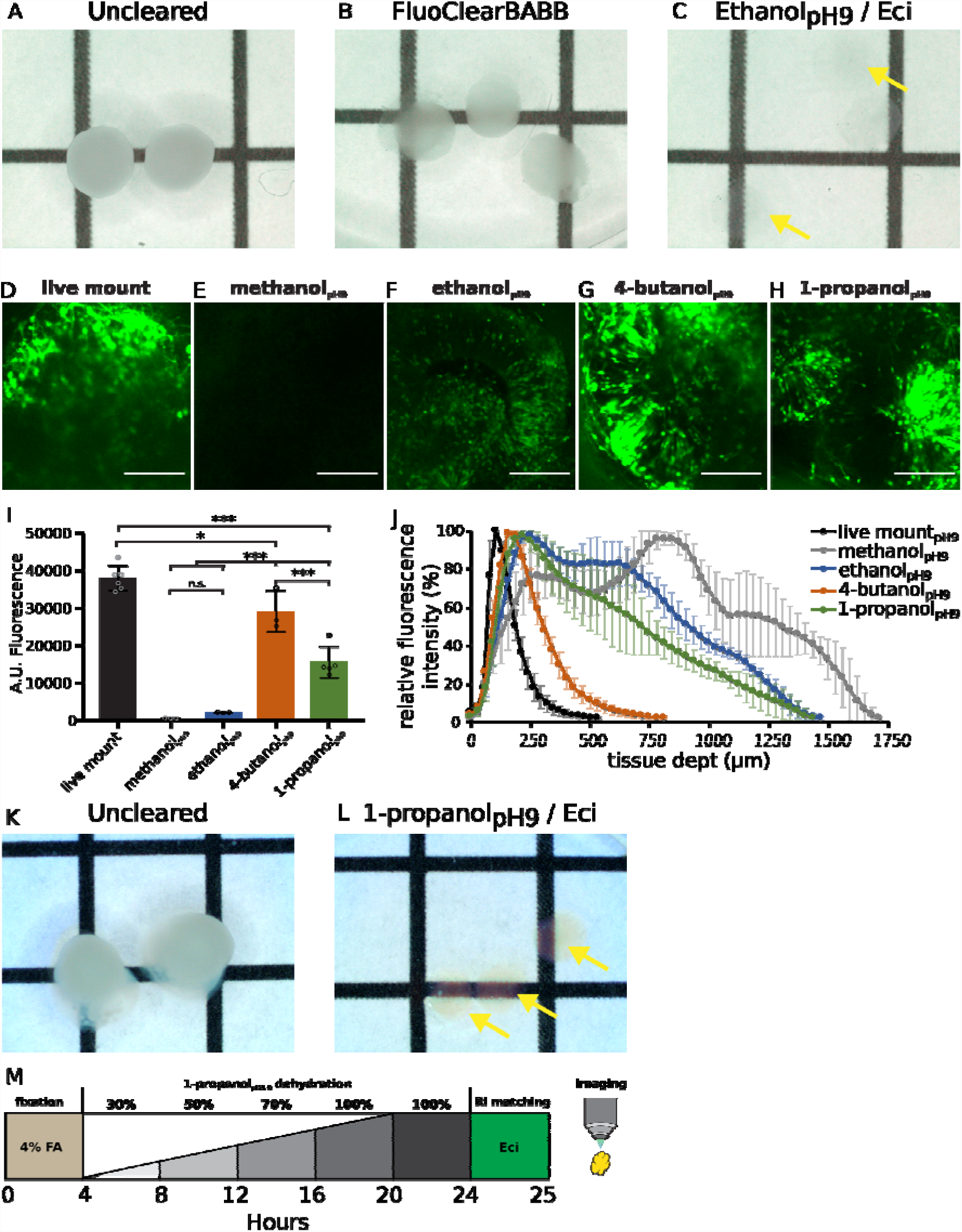
Eci clearing optimizations in cerebral organoids. Whole-mount recording of >100 day old cerebral organoids after fixation (A), FluoClearBABB (B) and ethanol _pH9_/Eci clearing. Yellow arrows mark organoids, 2 independent repetitions were performed. Dehydration agent dependent fluorescence after Eci mediated clearing (D-I). Comparison of live mounted organoids (D) and dehydration series (30%, 50%, 70%, 2x 100%, pH 9.0) of ethanol (E), methanol (F), 4-butanol (G) and 1-propanol (H) were performed on cerebral organoids and maximal fluorescence of Z stacks was quantified after Eci based clearing (I). N=3-6. Significance was calculated using ANOVA and post-hoc Turkey’s test. Quantification of tissue auto-fluorescence through alcohol- Eci cleared organoids as a measure of clearing efficiency (J). Uncleared organoids (K), are efficiently cleared (L) in as little as 25 hours including fixation, dehydration/delipidation, and refractive index matching. Scale bars: D-H 100 μm. Grid size A-C, K, L 5 mm.

To further validate 2Eci as an efficient method to clear cerebral organoids we recorded z-stacks through >100 days old cerebral organoids sparsely labeled with a population of CAG:GFP^+^-expressing cells. We found that 2Eci clearing allows for recordings throughout organoids of approximately 1400 μm thick (Fig. 2A, Movie 1), while the recording depth of uncleared cerebral organoids was approximately 100 μm into the tissue (Fig. 2B). Not only is imaging through an entire cerebral organoid possible, detailed morphological structures can be observed. In 80 day old cleared cerebral organoids, neural rosettes were readily observable in toto, with more mature neurons showing elaborate morphology engulfing the neuronal rosette (Fig. 2C). As with all dehydration-based methods, the organoids underwent a 30-40% reduction in diameter (Ertürk *et al.,* 2012) (Fig. S1B), but highly detailed morphological structures such as putative dendrites with dendritic spines and putative axons with boutons could still be observed (Fig. 2D-F). To examine whether GFP fluorescence is equally preserved throughout the organoid we whole mount labelled cerebral organoids with αGFP647 nanobody. We found consistent colocalization of GFP^+^ cells and the and the αGFP647 nanobody signal throughout the organoid (Fig. 2G-I, Fig. S1C). Taken together these data show that 2Eci is a viable method of clearing cerebral organoids and allows for detection of GFP signal within detailed morphological structures throughout the organoid.

**Figure 2:**
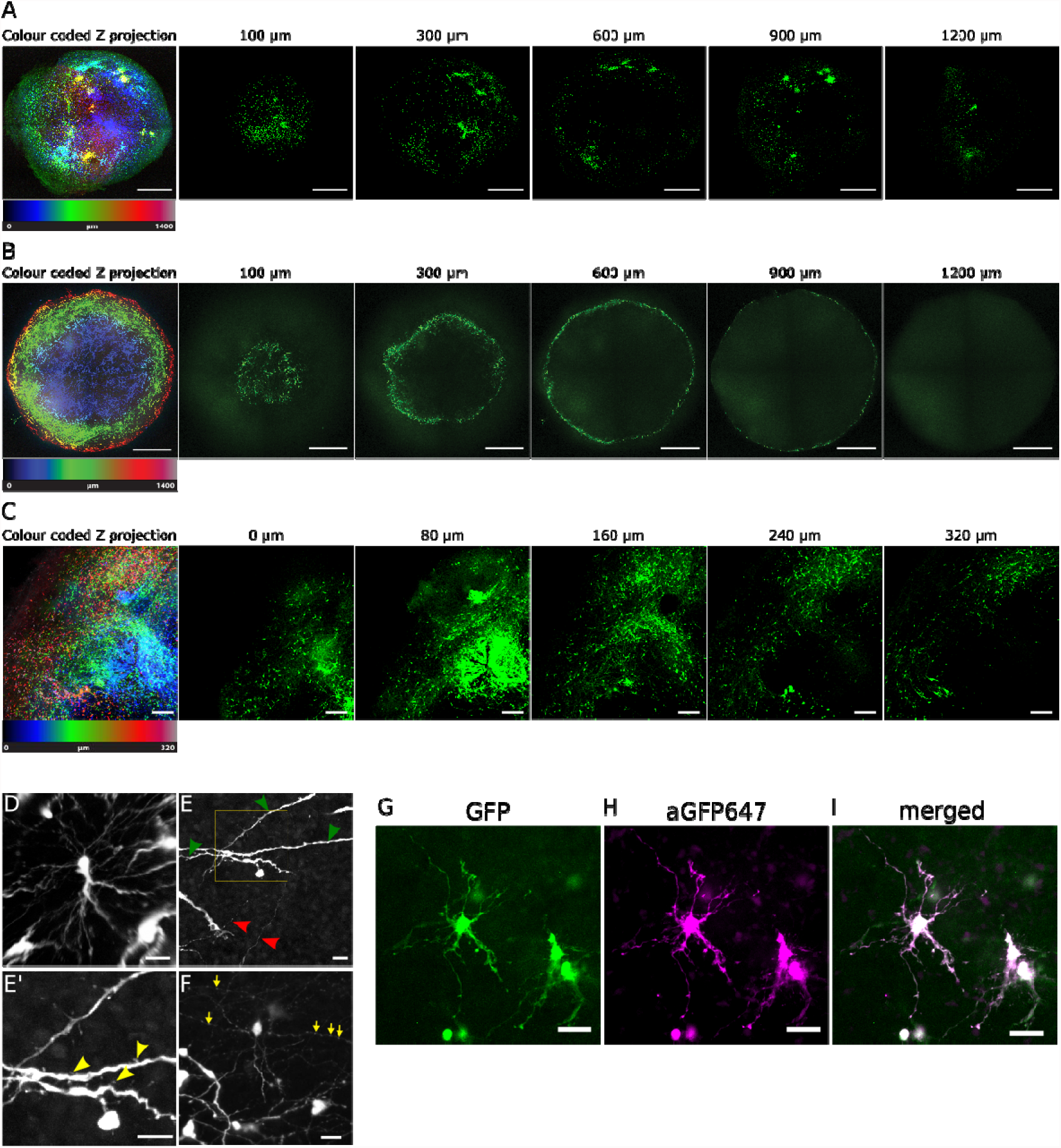
Characterization of 2Eci clearing. (A, B) Colour coded Z-projection and representative Z-slices of both 2Eci cleared (A) and uncleared (B) 3% GFP^+^ 80 day old cerebral organoids. 2Eci allows imaging through whole organoids. Spots of color aggregation depict aggregations of GFP^+^ neuronal stem cells in neuronal rosettes, whereas more mature neurons distribute more equally throughout the organoid. (C) Detailed morphology can be observed, including neuronal rosettes and more mature neurons in a day 90 old cerebral organoid. (D-F) 3D reconstruction of multiple neurons in 80 day old cerebral organoids. (D) Cellular details such as cell body shape and neurites can be observed using 20x objectives and 2x lens switch. (E) Putative dendrites (green arrowheads) and axons (red arrowheads) are maintained after clearing. (E’) Magnified view (yellow box) reveals putative dendritic spines (yellow arrowheads). (F) putative Boutons (yellow arrows) can be identified (G-I) αGFP647 accurately labels GFP fluorescence in single neurons. Scale bars: A-C 500 μm, D-F 10 μm, G-I 20 μm.

Any method that is to be used in a high-throughput approach should take cost and robustness into account. In order to interrogate the robustness of 2Eci while also significantly reducing cost, we interrogated the clearing efficiency of 2Eci in a large variety of different organisms and substituted the original 99% Ethyl cinnamate for the up to seven-fold cheaper 98% ethyl cinnamate. We first attempted to clear the hindlimb of a CAG:EGFP-expressing transgenic *Xenopus laevis.* The CAG:GFP transgene is ubiquitously expressed with highest levels of expression in the musculature. The hindlimb is a thick tissue but after removal of the pigmented skin the limbs could be cleared using Ethyl cinnamate so that the ubiquitous GFP fluorescence was observed throughout the limb (Fig. 3A-C, Movie 2).

**Figure 3:**
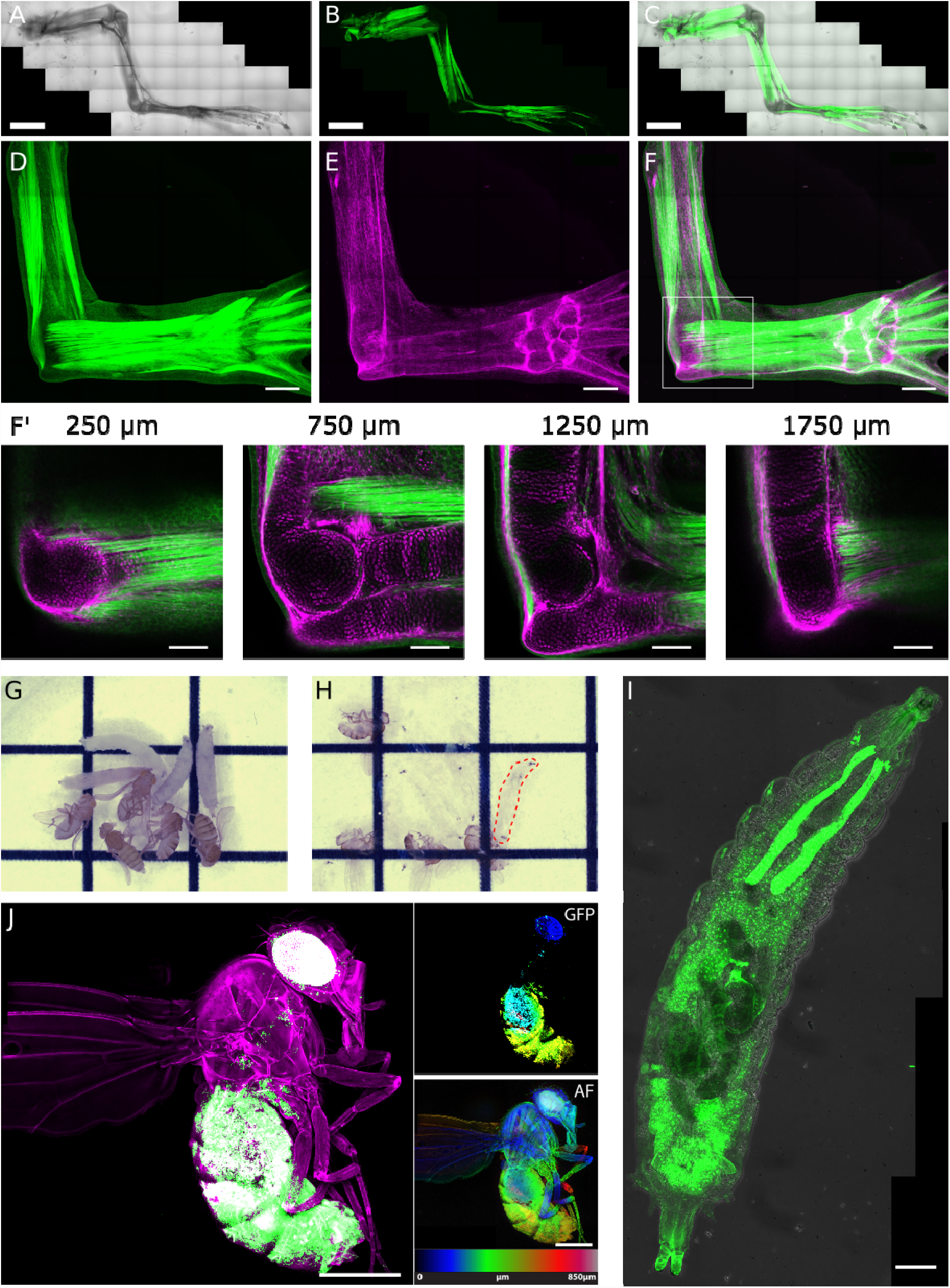
A large variety of tissues and animals are efficiently cleared using 2Eci while preserving signal. (A-C) CAG-GFP Xenopus laevis hind limbs are efficiently cleared using 2Eci. (A) brightfield imaged of cleared Xenopus laevis hind limb, (B) GFP is especially prevalent in the musculature, (C) merged view. (D-F) Both GFP (D, green), and mCherry (E, magenta) fluorescence is preserved after 2Eci clearing of Prrx1:ER-Cre-ER CAGGs:LP-GFP-LP-mCherry double transgenic axolotl. Detailed morphology can be observed throughout the limb (F’) including loose connective tissue, skeletal elements and tendons. (G-H) Drosophila larvae and adult drosophila before and after 2Eci clearing. Red dashed outline: indication of one of eight cleared drosophila larvae in the image (H). (I) Maximum intensity projection of a Krp-GFP drosophila larvae. Krp-GFP^+^ salivary gland and fat tissue can be observed throughout the larvae. (J) Whole Krp-GFP drosophila virgin z-Projection. Green represents GFP fluorescence, magenta represents autofluorescence. Right panels: Color coded z-projections of both GFP and 568nm autofluorescence (AF). Scale bars: A-C 2 mm, D-F 500 μm, F’ 200 μm: I-J 500 μm. Grid size G-H: 5 mm

To further interrogate the effectiveness of 2Eci we explored alternative fluorophores beyond GFP. Using a newly established *Prrx1:ER-Cre-ER; CAGGs:LP-GFP-LP-mCherry* double transgenic axolotl we used tamoxifen to induce recombination of the *CAGGs:LP-GFP-LP-mCherry* cassette. Recombination during larval and limb bud stages results in an indelible mCherry labelling of connective tissues throughout the limb(Logan *et al.,* 2002). These limbs can be efficiently cleared using 2Eci, resulting in the preservation of both GFP and mCherry signal (Fig. 3D-F), which can be observed throughout the entire depth of the limb (Fig. 3F’). To further explore fluorophore survival upon clearing we investigated the potential of 2Eci in clearing Brainbow tissue. While the recent development of antigen specific fluorophores in Brainbow3 allows for antibody labelling, previous constructs in both the original Brainbow and Brainbow2 series do not have antigen specificity (Cai *et al.,* 2013). To test fluorescent protein preservation, Axolotl Brainbow2.1R (Currie *et al.,* 2016) was crossed to CAGGs:ER-Cre-ER-T2A-EGFP-nuc (Khattak *et al.,* 2013) and recombination was induced followed by 2Eci clearing. We found that all Brainbow fluorophores are preserved and remain spectrally distinct (Fig. S1D) suggesting that 2Eci can also be used as a general tool for already existing Brainbow and Brainbow2 without having to rely on antibody labelling. A complete overview of tested fluorophores is provided in Table 2. To further challenge 2Eci, we cleared adult zebrafish, as they provide a unique challenge due to their size, scales and 3 types of pigment (xanthophores, melanophores, and iridophores). We found that while efficient clearing is achieved simply by increasing the time for dehydration and clearing steps (Fig. S2A-B) the silvery iridophores and black melanophores persist. Surprisingly the yellow xanthophores are efficiently cleared. The reason underlying this difference in pigment clearing is something we currently do not understand, but can be overcome using pigmentation mutants such as Nacre(Lister *et al.,* 1999) (Fig. S2C-D).

**Table 2:**
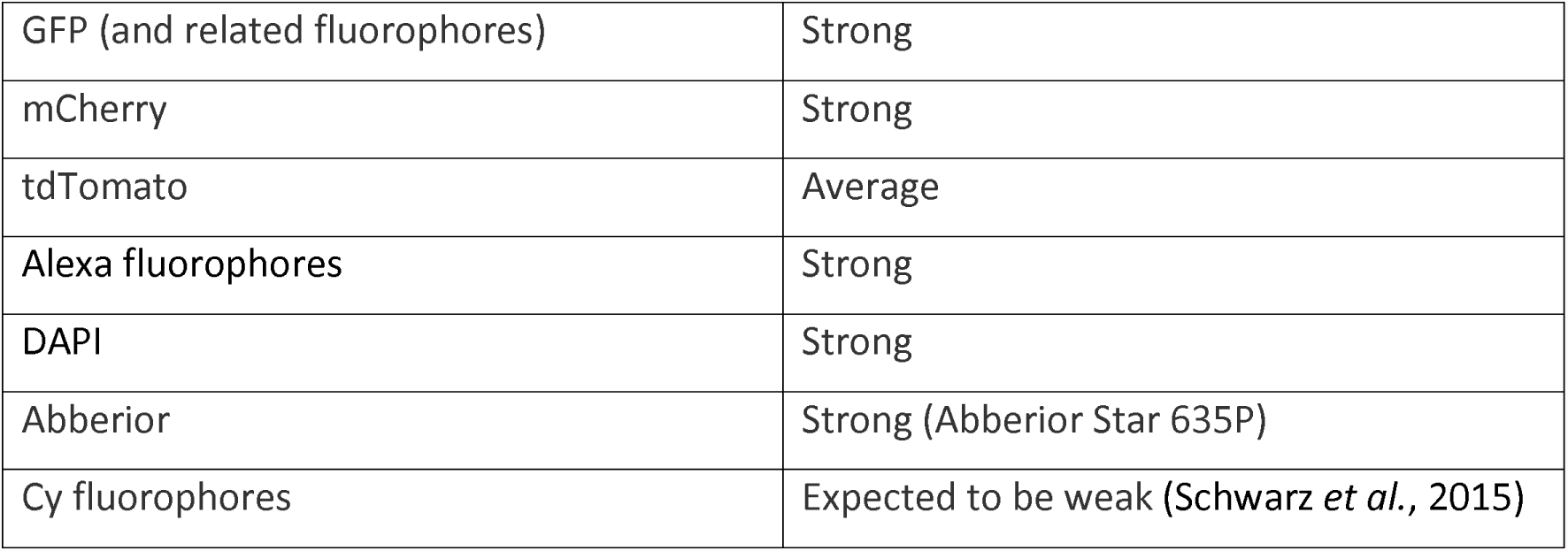
Overview of fluorescent probes or fluorophores compatibility with 2Eci clearing.

In the cerebral organoids, 2Eci surpassed FluoClearBABB in clearing effectiveness (Fig. 1B-C). We further investigated ethyl cinnamate as a viable clearing strategy after whole mount in-situ hybridization in amphibians, and in the context of adult *Drosophila melanogaster.* After WISH, amphibian embryos are commonly dehydrated using methanol and subsequently cleared using BABB (Saint-Jeannet, 2017). Adult *Drosophila melanogaster* were also previously shown to be efficiently cleared using BABB (McGurk *et al.,* 2007). We found Ethyl cinnamate to provide an efficient and non-toxic alternative in clearing of axolotl embryos after WISH (Fig. S1E). We thus expect embryos from other species that are also commonly cleared with BABB after WISH also to be efficiently cleared using Eci.

To test if adult *Drosophila melanogaster* can be cleared using Ethyl cinnamate, we used 2Eci to clear Krp:GFP transgenic drosophila. Prior to dehydration, we performed a CCD digest to digest parts of the exoskeleton and increase permeabilization (Manning and Doe, 2016). We found 2Eci to efficiently clear both larvae and adult drosophila (Fig. 3G-I), while preserving GFP expression (Fig. 3J-I). For adult drosophila, autofluorescence at 488 nm and 568 nm excitation was comparable. We therefore used the 568 nm channel to perform morphological reconstruction of drosophila and its inner organs and also subtract the auto fluorescent background of the 488 nm recording to visualize a GFP-specific signal. With this, whole fly reconstructions are possible while retaining the cellular resolution of Krp-GFP cells (Fig. 3J and Movie 3).

### 2Eci clearing can combine fluorescent proteins with antibody staining

Comparison of fluorescent protein expression with traditional immunofluorescent antibody staining is a workhorse method for characterizing cell types in complex tissues such as the limb. To interrogate if traditional two-step antibody labelling and fluorescence protein detection can be combined in 2Eci clearing, we used a combination of anti-Prrx1 antibody staining and Cre mediated lineage labelling in our transgenic *Col1a2:ER-Cre-ER; CAGGs:LP-GFP-LP-mCherry* reporter animals. Prrx1 is a broad marker of limb connective tissue while Col1a2 is a known marker of dermal fibroblasts and the skeletal lineage. Processing of limbs from the reporter transgenic showed that Col1a2-expressing cells localize to the skeletal, tendon and dermis. Antibody staining for PRRX1^+^ labeled cells in dermis peri-skeleton cells, and muscle interstitium with low intensity signal in the skeletal lineage (Fig. 4). This experiment highlights the heterogeneous nature of connective tissue. More importantly it further highlights utility of this rapid clearing protocol for combining fluorescent protein with immunofluorescence visualization to analyze the complex 3D morphologies of heterogeneous tissues.

**Figure 4:**
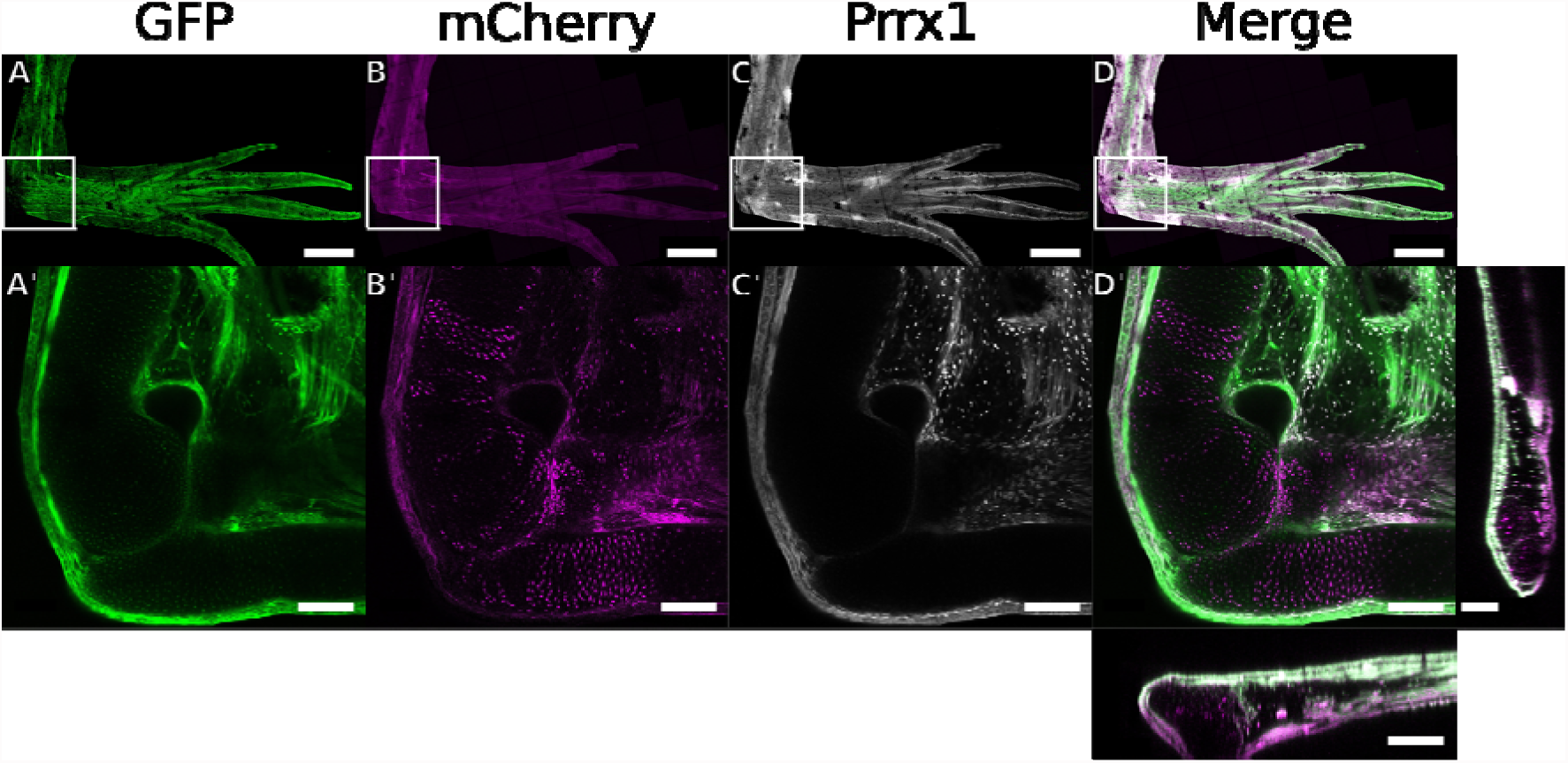
Heterogeneous nature of connective tissue as revealed by combining fluorophores and antibody staining in 2Eci cleared axolotl limbs. Upon Cre activity GFP^+^ cells (A), convert to an indelible mCherry^+^ labelling (B). Antibody staining (C) can be effectively combined with this resulting in a 3-channel image which highlights the complex heterogeneous nature of connective tissue (D). A-D are maximum intensity projections of the entire limb. The bounding box marks the elbow. Single slice recordings of the elbow are shown in A’-D’, D’ includes orthogonal views of the z-stack. Scale bars: A-D 1000 μm, A’-D’ 250 μm.

### Imaging considerations

Since Ethyl cinnamate is a non-toxic compound as opposed to BABB it can be used on microscopes in multi-user microscope facilities. However, there is a current lack of commercially available lenses optimized for Ethyl cinnamate both with regard to refractive index matching and immersion compatibility. Such lenses would ideally be applied in the context of high resolution light sheet microscopy. We opted instead to optimize deployment on commonly available inverted imaging platforms using low magnification(≤20x), low NA (<0.8) air objectives with long working distances. Such lenses provide a large field of view while reducing the effect of refractive index mismatching. This approach also prevents the lenses from coming into direct contact with Eci. While Eci is a non-toxic compound it is still a mild organic solvent and might attack insulation rings of objectives or imaging chambers. Thus, we set out to identify suitable commercially available mounting chambers for inverted imaging (table 3). From all the dishes that were tested, we identified the IBIDI μDish 35mm ibiTreat (cat.#81156) and Ibidi 35mm Glass Bottom (cat.#81158) as compatible with Ethyl cinnamate, being resistant to Ethyl cinnamate for at least several months. However, we recommend to store samples in air tight containers such as Falcon tubes or VOA glass vials filled with Ethyl cinnamate, as prolonged exposure to the air can result oxidation and mild declearing over time (data not shown). Samples which are difficult to orientate can be fixated in place by mounting samples in a 1% phytagel block prior to dehydration. After Clearing phytagel blocks can be mounted directly into the microscope dish using a small amount of super glue. Taken together this mounting and imaging approach should provide a reliable method which can be easily deployed in high throughput approaches using multi-user microscope facilities

**Table 3:** Overview of imaging dishes tested for compatibility with 2Eci clearing.

|  |  |
| --- | --- |
| Ibidi μSlide uncoated | Bottom falls out instantly (< 10 minutes) |
| Ibidi μSlide ibiTreat | Bottom falls out instantly (< 10 minutes) |
| **Ibidi μDish 35 mm high ibiTreat (81156)** | **Long term storage possible (>2 month)** |
| **Ibidi μDish 35mm high, Glass Bottom (81158)** | **Long term storage possible (>2 month)** |
| Ibidi μDish 35 mm high uncoated (81151) | Single recordings, gradually dissolves plastic |
| NUNC Lab-Tek (155361) | Single recordings, gradually dissolves plastic |
| NUNC Lab-Tek II (154453) | Single recordings, gradually dissolves plastic |
| Univer-slide (Alessandri *et al.*, 2017) | Long term storage possible |
| Lab-Tek II Chambered #1.5 Coverglass System (155409) 8 well chamber | Single recordings, gradually dissolves plastic |
| Lab-Tek II Chambered Coverglass System (154941) 8 well chamber | Single recordings, gradually dissolves plastic |
| Lab Tek chambered borsilicate coverglas 4 chamber (155383) | Single recordings, gradually dissolves plastic |
| Sarstedt 4-well auf Lumox detachable (94.6150.401) | Single recordings, gradually dissolves plastic |

We conclude that 2Eci is a broadly applicable clearing method that combines the rapid and broad applicability of dehydration-based methods, and the non-toxic nature and preservation of fluorescent proteins of aqueous clearing methods. This method preserves both fluorescent proteins while being also compatible with antibody staining. 2Eci clearing was shown to be effective in a large range of cases either matching or surpassing the efficacy of BABB. 2Eci is a simple, robust and cost-effective clearing method which should see use in high-throughput organoid screening approaches, among others.

## Acknowledgements

The authors acknowledge Francois Bonnay, Joshua Bagley, Tzi-Yang Lin, and Andrea Pauli for the contribution of tissues and cell lines, Tobias Müller for discussions, and the IMP BioOptics for availability and maintenance of microscopes.

## Competing interests

The authors declare no competing interests.

## Funding

WM is supported by a Lise Meitner fellowship (M2444); EMT is supported by ERC Advanced Grant, DFG grant TA 274/13-1, TA 274/3-3; JK is supported by the Austrian Academy of Sciences, the Austrian Science Fund (grants I_1281-B19 and Z-153-B09) and an advanced grant from the European Research Council.

## Movies

Movie 1

3D reconstruction of a 2Eci cleared 80-day old CAG-GFP cerebral organoid.

Movie 2

3D reconstruction of a 2Eci cleared *Xenopus laevis* CAG-GFP hind limb.

Movie 3

3D reconstruction of a 2Eci cleared Krp-GFP adult *Drosophila melanogaster.* Magenta labels auto-fluorescence, green marks Krp-GFP expression.

## Materials and Methods

### Growth of cerebral organoids

Organoids have been grown as described previously(Lancaster *et al.,* 2013) using feeder free H9 human embryonic stem cells (hES) from WiCell with a verified normal karyotype and contamination free. Cells were cultured in in mTESR1 (Cat.85850) and initial EB formation for the first 5 days was performed in mTESR1. For sparse labeling of organoids, 1-3% of the initial 9000 cells were replaced with feeder free H9 with a CAG-GFP insertion into AAVS1 (Bagley *et al.,* 2017) for initial EB formation.

### Animal husbandry and handling

Axolotl were maintained on a 12-h light/ 12-h dark cycle at 18-20°C(Khattak *et al.,* 2014). Prior to amputation or tissue collection animals were anaesthetized in 0.03 % benzocaine and injected subcutaneously with 38mg/Kg buprenorphine. 0.1% benzocaine was used for terminal experiments and euthanasia of axolotl. The work was performed under an approved license from the Magistrat der Stadt Wien (GZ: 9418/2017/12).

TLAB and Nacre(Lister *et al.,* 1999) zebrafish were maintained on a 14-h light/ 10-h dark cycle at 28°C according to standard procedures(Westerfield, 1995). TLAB fish, generated by crossing zebrafish AB and the natural variant TL (Tübingen/Tüpfel Longfin) stocks, served as wild-type zebrafish. All fish experiments were conducted according to Austrian and European guidelines for animal research and approved by local Austrian authorities (animal protocol BMGF-76110/0017-II/B/16C/2017).

Xenopus were maintained on a 12-h light/ 12-h dark cycle at 20°C. Prior to amputation or tissue collection animals were anaesthetized in 0.01% MS222 and injected subcutaneously with 38mg/Kg buprenorphine. The work is performed under an approved license from the Magistrat der Stadt Wien (GZ: 852533/2016/20).

The following *Drosophila melanogaster* stock was used in this study: w; L^2^ Pin^1^/CyO, P{GAL4-Kr.C}DC3, P{UAS-GFP.S65T}DC7 (Bloomington #5194), a larval fat-body-expressing GFP reporter. Wandering third instar larvae and young virgin females still displaying larval fat-body were used as material for clearing.

### Clearing of cerebral organoids

Organoids were fixed in 4% PFA for 4h at RT or at 4°C over night and transferred sequentially into a dehydration series of 30%, 50%, 70% and 2x 100% 1-Propanol (99%; Sigma Cat.W292818, 99.7%: Sigma Cat. 27944): 1xPBS solution pH adjusted to 9.0 – 9.5 using trimethylamine (Sigma Cat. T0886). For comparison of dehydration agents, 1-Propanol was exchanged with 4-Butanol, Ethanol or Methanol respectively. Dehydration was performed at 4°C on a gyratory rocker for at least 4h per dehydration step in 50ml Falcon tubes containing 45ml dehydration agent.

Subsequently, organoids were transferred in a 50ml tube with at least 25ml ECi (≥98%: Sigma Cat. W243000, 99%: Sigma Cat. 112372) and incubated on a gyratory rocker at room temperature for at least one hour before recording. Samples were stored in light-protected and air-sealed containers. Recordings were acquired over the following days.

### IHC of cerebral organoids

IHC was optionally performed after 4% PFA fixation. In brief, organoids were washed in PBS for 10min to remove residual PFA. Subsequently, the organoids were transferred in permeabilization/blocking solution (0.3% TX100, 5% BSA, 0.05% NaN_3_ in PBS) over night. For antibody staining, a GFP booster in far red (Atto647, Chromotek GBA647n) at 1:50 dilution was used in antibody staining solution (0.1%TX100, 5%BSA, 0.05% NaN_3_). IHC was performed at 37°C for 2 days. Subsequently, organoids were washed in PBS-T (PBS and 0.1% TX100) for one day. The organoids were then fixed in 4% PFA and used for clearing as described.

### Clearing and recording of drosophila

Drosophila and drosophila larvae were used for clearing experiments. Drosophila were transferred into 30% EtOH to remove the hydrophobic fatty lipid layer from the exoskeleton for 5-10 minutes. Subsequently, drosophila were briefly bleached using DanKlorix, a commercially available bleach solution (Colgate-Palmolive). To increase transparency of the exoskeleton, a CCD (Chitinase-chymotrypsin-DMSO buffer) digest was performed(Manning and Doe, 2016). Drosophila were then subsequently fixed in 4% PFA in PBS for 4h at RT or at 4°C over night. Clearing was performed as for cerebral organoids, however the incubation in Ethyl cinnamate was extended to 3+ hours. For adult fly recordings, the GFP fluorescence (488nm excitation, Filter: 525/50) as well as the autofluorescence of 561nm (561nm excitation, Filter: 609/54) was recorded and the autofluorescence of the 561 channel was subtracted from the GFP recording. To ensure autofluorescence specificity in the subtraction process, the 562 channel was recorded with autofluorescence intensity levels below the levels of GFP autofluorescence. Alternatively, the intensity of the 561 channel was modulated to achieve levels slightly below GFP autofluorescence levels. We did not find significant differences between both 488nm/525 and 561nm/609 autofluorescence. Additionally to autofluorescence correction, the 561/609 recording was used to reconstruct morphological details of adult drosophila.

### Clearing of axolotl tissue

Axolotl tissue was harvested as previously described(Roensch *et al.,* 2013), briefly axolotl tissue fixed at 4°C over night in 1x MEMFA (0.1M MOPS pH 7.4, 2mM EGTA,s 1mM Mg SO4 × 7H2O and 3.7% formaldehyde), washed in PBS and cleared as for cerebral organoids, however dehydration and Ethyl cinnamate incubation steps were increased to 12 hours.

### IHC of whole mount axolotl tissue

IHC was performed optionally after MEMFA fixation. Tissue was washed in PBS at RT for 2x 1 hour, followed by a 3x 2 hour PBS-T (0.3% triton) wash. Blocking was performed at 37°C overnight in PBS-T supplemented with 5% bovine serum. Staining was performed at 37°C for 48 hours in PBS-T supplemented with 5% bovine serum and axolotl prrx1 antibody (Ocaña *et al.,* 2017). Tissue was again washed and blocked, followed by staining for secondary antibodyTissue was extensively washed in PBS after staining and processed for clearing.

### Transgenesis and lineage tracing in axolotl

To label connective tissue populations in in axolotl the newly generated lines of Col1a2:ER-Cre-ER, and Prrx1:ER-Cre-ER were onto the already existing CAGGs:LP-EGFP-LP-mCherry(Khattak *et al.,* 2013). To generate the Prrx1 and Col1a2 lines, the Prrx1 enhancer/promoter (Logan *et al.,* 2002)(a kind gift from Malcolm Logan) and the Col1A2 promoter(Bou-Gharios *et al.,* 1996) (a kind gift from George Bou-Gharious) was cloned at the 5’ end of TFPnls-T2A-ERT2-Cre-ERT2 (ER-Cre-ER) cassette with flanking SceI sites. Transgenesis was performed as previously described(Khattak *et al*., 2014). 4-OHT treatment is done as described previously(Khattak *et al*., 2014). Briefly, 3 cm long double transgenic animals were treated with 2 μM 4-Hydroxy Tamoxifen (4-OHT) by bathing over night. Tissue was collected 2 weeks post treatment.

### Whole mount In-situ hybridizations of axolotl

Chromogenic in-situ hybridizations were performed as previously described (Cerny *et al.,* 2004) using stage 35 axolotl. Probes for GFAP(Rodrigo Albors *et al.,* 2015) were generated as previously described. After staining and dehydration axolotl embryos were incubated 15-30 minutes in 98% Ethyl Cinnamate to clear the tissue.

### Zebrafish fixation and clearing

Zebrafish were collected and fixed in 4% PFA using standard procedures(Westerfield, 1995). Clearing was performed as described for organoids however dehydration and ethyl cinnamate incubation steps were increased to 12 hours. Additionally, the swim bladder was pierced and allowed to fill with ethyl cinnamate.

### Xenopus fixation and clearing

Xenopus hind limbs were collected as for axolotl. The pigmented skin was carefully removed from the hind limbs using forceps. Hind limbs were cleared as described for cerebral organoids however dehydration and ethyl cinnamate incubation steps were increased to 12 hours.

### Microscopy

Organoid and drosophila recordings were performed on a Yokogawa W1 spinning disk confocal microscope (VisiScope, Visitron Systems GmbH, Puchheim, Germany) controlled with VisiView Software (Visitron) and mounted on the Eclipse Ti-E microscope (Nikon, Nikon Instruments BV). Recordings were performed with a 10x/0.45 CFI plan Apo Lambda, 20x/0.75 CFI plan Apo lambda or CFI plan Apo lambda 40x/1.4 oil (all: Nikon, Nikon Instruments BV) objectives with a sCMOS camera (PCO edge 4.2m, PCO AG) or an EMCCD camera (Andor Ixon Ultra 888). For stitching, the stitching plugin in Fiji (based on ImageJ 1.51k) was used. For 3D reconstructions, the freeware Icy (Version 1.9.5.1) was used. Axolotl and Xenopus recordings were collected on an inverted Zeiss LSM780 equipped with a 10x/0.3 EC plan-neofluar objective (Carl Zeiss Microscopy GmbH, Germany). Zen 2.3 SP1(black) (64 bit) was used for image acquisition and automatic stitching of images. Image preparation was performed using FIJI (based on ImageJ 1.51k) and Inkscape 0.91 (www.inkscape.org). Adult zebrafish and intact axolotl recordings were acquired using a Zeiss Lumar stereomicroscope (Carl Zeiss Microscopy GmbH, Germany) equipped with Spot Pursuit-XS monochrome and Spot Insight color cameras (SPOT Imaging USA).

### Statistics

One way-ANOVA and post-hoc Tukey’s test were performed to determine significance between groups.

## References

Alessandri, K., Andrique, L., Feyeux, M., Bikfalvi, A., Nassoy, P. and Recher, G. (2017) ‘Allin-one 3D printed microscopy chamber for multidimensional imaging, the UniverSlide.’, Scientific reports, 7, p. 42378. doi: 10.1038/srep42378.

Bagley, J. A., Reumann, D., Bian, S., Lévi-Strauss, J. and Knoblich, J. A. (2017) ‘Fused cerebral organoids model interactions between brain regions’, Nature Methods, 14(7), pp. 743–751. doi: 10.1038/nmeth.4304.

Belle, M., Godefroy, D., Couly, G., Malone, S. A., Collier, F., Giacobini, P. and Chédotal, A. (2017) ‘Tridimensional Visualization and Analysis of Early Human Development.’, Cell, 169(1), pp. 161–173.e12. doi: 10.1016/j.cell.2017.03.008.

Belle, M., Godefroy, D., Dominici, C., Heitz-Marchaland, C., Zelina, P., Hellal, F., Bradke, F. and Chédotal, A. (2014) ‘A simple method for 3D analysis of immunolabeled axonal tracts in a transparent nervous system.’, CellReports, 9(4), pp. 1191–1201. doi: 10.1016/j.celrep.2014.10.037.

Bou-Gharios, G., Garret, L. A., Rossert, J., Niederreither, K., Eberspaecher, H., Smith, C., Black, C. and de Crombrugghe, B. (1996) ‘A potent far-upstream enhancer in the mouse pro alpha 2(I) collagen gene regulates expression of reporter genes in transgenic mice’, The Journal of Cell Biology, 134(5), pp. 1333–1344. doi: 10.1083/jcb.134.5.1333.

Cai, D., Cohen, K. B., Luo, T., Lichtman, J. W. and Sanes, J. R. (2013) ‘Improved tools for the Brainbow toolbox.’, Nature Methods, 10(6), pp. 540–547. doi: 10.1038/nmeth.2450.

Cerny, R., Lwigale, P., Ericsson, R., Meulemans, D., Epperlein, H. H. and Bronner-Fraser, M. (2004) ‘Developmental origins and evolution of jaws: new interpretation of “maxillary” and “mandibular”.’, Developmental Biology, 276(1), pp. 225–236. doi: 10.1016/j.ydbio.2004.08.046.

Chung, K., Wallace, J., Kim, S.-Y., Kalyanasundaram, S., Andalman, A. S., Davidson, T. J., Mirzabekov, J. J., Zalocusky, K. A., Mattis, J., Denisin, A. K., Pak, S., Bernstein, H., Ramakrishnan, C., Grosenick, L., Gradinaru, V. and Deisseroth, K. (2013) ‘Structural and molecular interrogation of intact biological systems’, Nature. Nature Publishing Group, 497(7449), pp. 332–337. doi: 10.1038/nature12107.

Currie, J. D., Kawaguchi, A., Traspas, R. M., Schuez, M., Chara, O. and Tanaka, E. M. (2016) ‘Live Imaging of Axolotl Digit Regeneration Reveals Spatiotemporal Choreography of Diverse Connective Tissue Progenitor Pools.’, Developmental Cell, 39(4), pp. 411–423. doi: 10.1016/j.devcel.2016.10.013.

Economo, M. N., Clack, N. G., Lavis, L. D., Gerfen, C. R., Svoboda, K., Myers, E. W. and Chandrashekar, J. (2016) ‘A platform for brain-wide imaging and reconstruction of individual neurons’. eLife Sciences Publications Limited, 5, p. e10566. doi: 10.7554/eLife.10566.

Ertürk, A., Becker, K., Jährling, N., Mauch, C. P., Hojer, C. D., Egen, J. G., Hellal, F., Bradke, F., Sheng, M. and Dodt, H.-U. (2012) ‘Three-dimensional imaging of solvent-cleared organs using 3DISCO.’, Nature Protocols. Nature Publishing Group, 7(11), pp. 1983–1995. doi: 10.1038/nprot.2012.119.

Ke, M.-T., Fujimoto, S. and Imai, T. (2013) ‘SeeDB: a simple and morphology-preserving optical clearing agent for neuronal circuit reconstruction’, Nature Neuroscience. Nature Publishing Group, 16(8), pp. 1154–1161. doi: 10.1038/nn.3447.

Khattak, S., Murawala, P., Andreas, H., Kappert, V., Schuez, M., Sandoval-Guzmán, T., Crawford, K. and Tanaka, E. M. (2014) ‘Optimized axolotl (Ambystoma mexicanum) husbandry, breeding, metamorphosis, transgenesis and tamoxifen-mediated recombination.’, Nature Protocols, 9(3), pp. 529–540. doi: 10.1038/nprot.2014.040.

Khattak, S., Schuez, M., Richter, T., Knapp, D., Haigo, S. L., Sandoval-Guzmán, T., Hradlikova, K., Duemmler, A., Kerney, R. and Tanaka, E. M. (2013) ‘Germline transgenic methods for tracking cells and testing gene function during regeneration in the axolotl.’, Stem cell reports, 1(1), pp. 90–103. doi: 10.1016/j.stemcr.2013.03.002.

Klingberg, A., Hasenberg, A., Ludwig-Portugall, I., Medyukhina, A., Männ, L., Brenzel, A., Engel, D. R., Figge, M. T., Kurts, C. and Gunzer, M. (2017) ‘Fully Automated Evaluation of Total Glomerular Number and Capillary Tuft Size in Nephritic Kidneys Using Lightsheet Microscopy.’, Journal of the American Society of Nephrology : JASN. American Society of Nephrology, 28(2), pp. 452–459. doi: 10.1681/ASN.2016020232.

Lancaster, M. A., Renner, M., Martin, C.-A., Wenzel, D., Bicknell, L. S., Hurles, M. E., Homfray, T., Penninger, J. M., Jackson, A. P. and Knoblich, J. A. (2013) ‘Cerebral organoids model human brain development and microcephaly.’, Nature. Nature Publishing Group, 501(7467), pp. 373–379. doi: 10.1038/nature12517.

Lister, J. A., Robertson, C. P., Lepage, T., Johnson, S. L. and Raible, D. W. (1999) ‘nacre encodes a zebrafish microphthalmia-related protein that regulates neural-crest-derived pigment cell fate’, Development (Cambridge, England), 126(17), pp. 3757–3767. doi: 10.1074/jbc.273.31.19560.

Logan, M., Martin, J. F., Nagy, A., Lobe, C., Olson, E. N. and Tabin, C. J. (2002) ‘Expression of Cre recombinase in the developing mouse limb bud driven by aPrxl enhancer’, genesis, 33(2), pp. 77–80. doi: 10.1002/gene.10092.

Manning, L. and Doe, C. Q. (2016) ‘Immunofluorescent antibody staining of intact Drosophila larvae’, Nature Protocols, 12(1), pp. 1–14. doi: 10.1038/nprot.2016.162.

McGurk, L., Morrison, H., Keegan, L. P., Sharpe, J. and O’Connell, M. A. (2007) ‘Three-Dimensional Imaging of Drosophila melanogaster’, PLoS ONE. Edited by S. Rutherford, 2(9), pp. e834–6. doi: 10.1371/journal.pone.0000834.

Ocaña, O. H., Coskun, H., Minguillón, C., Murawala, P., Tanaka, E. M., Galcerán, J., Muñoz-Chápuli, R. and Nieto, M. A. (2017) ‘A right-handed signalling pathway drives heart looping in vertebrates.’, Nature. Nature Publishing Group, 549(7670), pp. 86–90. doi: 10.1038/nature23454.

Rodrigo Albors, A., Tazaki, A., Rost, F., Nowoshilow, S., Chara, O. and Tanaka, E. M. (2015) ’Planar cell polarity-mediated induction of neural stem cell expansion during axolotl spinal cord regeneration.’, 4, p. e10230. doi: 10.7554/eLife.10230.

Roensch, K., Tazaki, A., Chara, O. and Tanaka, E. M. (2013) ‘Progressive Specification Rather than Intercalation of Segments During Limb Regeneration’, Science, 342(6164), pp. 1375–1379. doi: 10.1126/science.1241796.

Saint-Jeannet, J.-P. (2017) ‘Whole-Mount In Situ Hybridization of Xenopus Embryos.’, Cold Spring Harbor Protocols, 2017(12), p. pdb.prot097287. doi: 10.1101/pdb.prot097287.

Schwarz, M. K., Scherbarth, A., Sprengel, R., Engelhardt, J., Theer, P. and Giese, G. (2015) ‘Fluorescent-protein stabilization and high-resolution imaging of cleared, intact mouse brains.’, PLoS ONE, 10(5), p. e0124650. doi: 10.1371/journal.pone.0124650.

Westerfield, M. (1995) The zebrafish book: a guide for the laboratory use of zebrafish (Brachydanio rerio).

